# Detecting genomic regions enriched for reciprocal recombination in autism spectrum disorder

**DOI:** 10.64898/2026.05.26.727863

**Authors:** Catherine F. Mahoney, Michael Salter-Townshend, Darren J. Fitzpatrick, Denis C. Shields

## Abstract

Meiotic recombination is an important means of increasing genetic diversity by generating novel haplotypes in a population. Recombination separates linked loci extremely slowly in some regions, therefore genetic variants in high linkage disequilibrium may become co-adapted. Reciprocal recombination that separates co-adapted variants may generate a deleterious *de novo* haplotype that contributes to disease. We developed statistical methods to detect genomic regions of recombination excess in two different family-based study designs. We identified recombination in the Simons Simplex Collection in 273 simplex families with one child with autism spectrum disorder (ASD) and at least two unaffected children, in which recombinations can be mapped to the proband and contrasted with the recombination counts in unaffected siblings; and in 1,802 families with two children, where the number of recombinations identified can be contrasted with the expectation from a reference recombination map. Both strategies revealed a tail of low p-values for loci of interest that contrasted with the rest of the distribution. Permutation and bootstrap tests did not identify genome-wide primary findings in either cohort, but the most significant three-child cohort locus of recombination excess (between cadherin genes *CDH4* and *CDH26*) replicated in the two-child cohort (p=0.01). While this replication strategy was not defined *a priori*, five of the most recombination enriched bins identified candidate ASD genes (p=0.02; *WWOX, ADAMTS16, INSR, ADARB2, and HS6ST1)*. Since the six identified loci were not identified as regions of high *de novo* copy number variation in the study cohort and no CNVs were detected in any of the recombinant probands in the identified regions, they represent candidates for reciprocal recombinations generating unfavourable haplotypes for these genes. This study highlights a previously unidentified source of clinical genetic variability contributing to the molecular aetiology of ASD.

**AUTHOR SUMMARY:** Autism spectrum disorder (ASD) is a constellation of neurodevelopmental disabilities characterised by deficits in social communication and repetitive patterns of behaviour. While ASD is highly heritable, its genetic basis is complex and poorly understood. While some highly penetrant types of genetic variation have been identified, most people with ASD carry a large number of variants that each contribute a small amount to their overall phenotype. In addition to mutations in individual genes, changes in the configuration of genes along a chromosome may contribute to ASD. Here, we describe a method for identifying regions where such new configurations have occurred through recombination and attempt to find regions where such changes are more common in autistic children than in their non-autistic siblings. We explore recombination as a source of genetic variation contributing to autism, which has potential to inform clinicians in providing services to autistic people and their families.

## INTRODUCTION

Meiotic recombination is a vital contributor to genetic variation in human populations. The process of recombination serves to improve selection efficiency in a population by allowing for new combinations of advantageous variants on a single chromosome, and counteracting the otherwise irreversible accumulation of deleterious mutations via “Muller’s ratchet” (1–6). The rate of meiotic recombination varies over the human genome, resulting in segments of chromosomes where little variation is generated over many generations (6). There is evidence of co-adapted variants in regions of high linkage disequilibrium in *Drosophila melanogaster* and chum salmon (*Oncorhychus keta)*. It is therefore plausible that there exist co-adapted haplotype pairs in humans, possibly in recombination-poor genomic regions, such that a *de novo* recombination occurring between a co-adapted pair of variants can contribute to disease (7,8).

We examine whether there is an association between *de novo* recombination events and increased probability of autism spectrum disorder by comparing the locations of recombination hotspots in autistic children versus non-autistic children. One well studied mechanism by which recombination can alter the configuration of a genetic locus is via changes in copy number; numerous studies have identified an association between copy number variation and autism spectrum disorder (9–11). This study, in contrast, examines the contribution of reciprocal recombination. Reciprocal recombination refers to a recombination event in which the amount of genetic material is conserved without insertion, deletion, or change in copy number. The association of recombination and novel haplotypes with disease is a relatively new area of research. The most apparent mechanism by which recombination and the generation of *de novo* haplotypes might contribute to disease is through changes in the expression of particular alleles. Broadly speaking, several studies have found that purifying selection depletes haplotypes that are likely pathogenic, but selection efficiency varies across the genome according to effective population size and recombination rate (12–16). Recombination is more effective at removing deleterious mutations in areas where there is a high rate of recombination, and thus areas of low recombination can accumulate a relatively higher number of deleterious mutations (15).

Two studies have examined the role of haplotype configuration in the accumulation and expression of putatively pathogenic variants. Castel, *et al* examined the effect of regulatory modifiers of penetrance on disease risk in both cancer and autism. In disease-associated genes, rare pathogenic variants that were exclusive to affected patients were significantly enriched for haplotype configurations where the major allele was the lower-expressed variant. Among control-specific variants, no such association was observed (13). Harwood, *et al* further examined the effect of recombination rate on the expression of pathogenic variants. The authors found that among loci where an underexpressed derived SNP is on the same haplotype as an eQTL cis-regulatory variant, the concordant haplotypes that underexpress the pathogenic variant are more likely to be located in hotspots or areas of average recombination rates as compared to coldspots. This phenomenon supports the hypothesis that allele specific expression is more effective at mitigating the effect of pathogenic variants in areas of high recombination (14).

We introduce a number of novel methods to determine whether there are areas where enrichment for *de novo* recombination is associated with autism spectrum disorder (ASD). Autism spectrum disorder is highly relevant for the study of *de novo* recombination, as there are several examples of causal *de novo* mutation in autism, and novel haplotypes generated by recombination may be considered to be *de novo* variation (17). The analysis in this paper aims to identify recombination hotspots in autistic children versus their unaffected siblings. Each of the methods was developed with the aim of balancing sufficient power to detect enrichment while providing adequate resolution to examine the gene content of the hotspots. A number of specially devised resampling based statistical methods facilitate the estimation of the statistical power of each hotspot detection strategy and the control of the family-wise error rate in tests of statistical significance. To maximise power, the study focuses on hotspot detection in the combined set of recombinations from both parents, but includes subgroup analyses of maternal and paternal recombinations. The rationale in including the parental subgroup analyses was that the pattern of recombination differs dramatically between parental sexes and there are several well known examples of parent-of-origin effects in developmental disorders, such as maternally-derived supernumerary chromosome 15 and Prader-Willi/Angelman Syndromes (18,19).

## MATERIALS AND METHODS

Recombination excesses were investigated in aggregate as the primary endpoint, since the pooled recombination data had a greater sample size and therefore increased statistical power, and secondarily among maternally and paternally derived recombination events, since the recombination maps differ extensively during the meiotic production of male and female gametes. Intervals longer than 200kb were excluded from analysis, on the basis that extremely long intervals would not be informative for local hotspots given that areas of linkage disequilibrium typically do not extend further than approximately this distance. Such long intervals could also reflect large structural changes such as deletions, rather than reciprocal recombination events. An R/Bioconductor package called *inferRecom* was developed to define each recombination event, resolved to an interval between two informative SNPs.

Briefly, the method implemented in *inferRecom* for detecting homologous recombination compares child genotypes to each other at a subset of single nucleotide polymorphisms (SNPs) that are informative for recombination, as illustrated in Fig 1, adapted from the original paper by Chowdhury, *et al* that describes the method (20). An informative SNP occurs where one parent is homozygous for the SNP and the other parent is heterozygous. For instance, SNPs where the father is homozygous and the mother is heterozygous are informative for recombination in the children’s inherited maternal chromosome, as each child will have inherited the same alleles from the father, making it possible to determine which alleles each child has inherited identical-by-descent from the mother. Comparing two siblings, a switch from one maternal haplotype to another indicates a recombination in one of the two children between the two informative SNPs. Determining in which sibling the recombination occurred, however, requires a third sibling. A switch in the maternal haplotype in one sibling as compared to the others indicates a recombination event in that child’s chromosome. Given that recombination events are relatively rare, typically occurring between one and three times per chromosome, it is very unlikely that two siblings will have a recombination occur in the same interval between informative SNPs (20–22).

**Fig 1.**
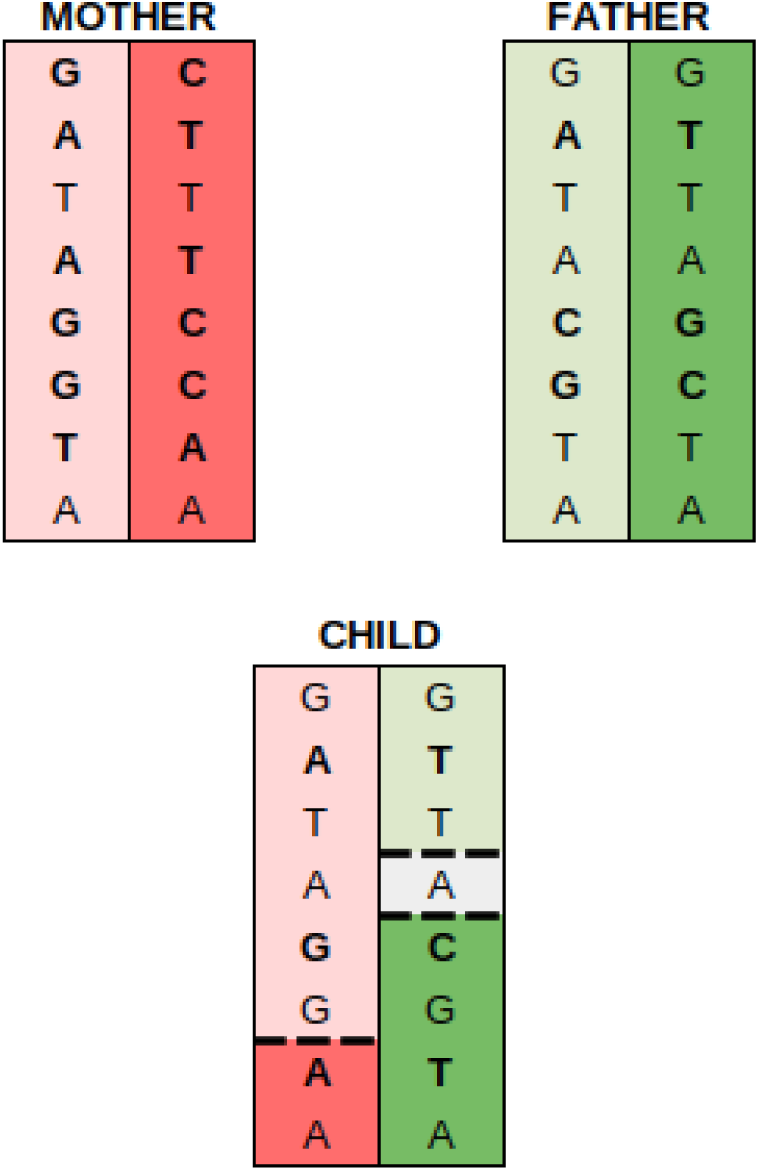
Recombination detection in phased chromosomes. For illustrative purposes, a child is shown that has two recombinations in a short interval, one on the maternally inherited chromosome and one on the paternally inherited. When haplotype phase is determined (through the presence of at least two other siblings identifying the two haplotypes within a parent), recombination can only be resolved to the nearest heterozygous parental SNP. In the maternal chromosome, the dashed line indicates the location of the recombination, which occurs between two heterozygous SNPs and is therefore known precisely. In contrast, the two dashed lines in the paternal chromosome indicate the two possible locations of the recombination, as there is a homozygous paternal SNP between the two heterozygous markers. See Chowdhury, *et al.* (20).

### Recombination hotpot analysis in fully informative pedigrees

Families with one autistic child and two or more non-autistic children are classed as ‘fully informative,’ as the presence of three or more children allows for the recombinant child in a nuclear family structure to be identified. In fully informative pedigrees, direct case-control comparison of the density of recombination intervals in autistic children versus non-autistic siblings is therefore possible. Genetic data from SSC v15.3, mapped to the hg19/GRCh37 reference genome, were used in this study. Quality control was performed in PLINK 1.9, beginning with filtering for Mendel errors and removal of any misattributed parent-child relationships to account for non-paternity or non-maternity events (23). Individuals and variants with a missingness rate exceeding 10% were excluded, the minimum minor allele frequency was set at 0.01, and SNPs deviating from Hardy-Weinberg equilibrium (chi-square p < .0001) were removed (24). Principal components analysis was applied to control for population stratification, which can introduce spurious associations in genome-wide studies due to systematic differences in allele frequencies between groups (25,26). The largest cluster, comprising approximately 90% of samples and corresponding largely to participants of European ancestry was retained, as it was the only group of sufficient size for analysis. Family structures were then filtered to include only those comprising exactly two parents, one affected child, and at least one unaffected child. Of the 2,587 families in the dataset, 273 three-child families and 1,802 two-child families passed both the PLINK and pedigree quality control steps and were retained for analysis. Given the limited sample size, it was necessary to devise a maximally powerful method for detecting and testing the enrichment of putatively enriched regions of *de novo* recombination, hereafter referred to as bins.

Two recombination hotspot bin definitions were explored. “Cluster binning” detected regions where a minimum number of case child recombination intervals were clustered with a maximum gap of one average gene length (15kb) between intervals. “Minimum depth binning” detected regions of the genome where a minimum number of recombination intervals overlap a single point, with the bin limits defined as the furthest extent of maximal recombination interval overlap, plus a 15kb buffer on either side. Both methods are illustrated in Fig 2.

**Fig 2.**
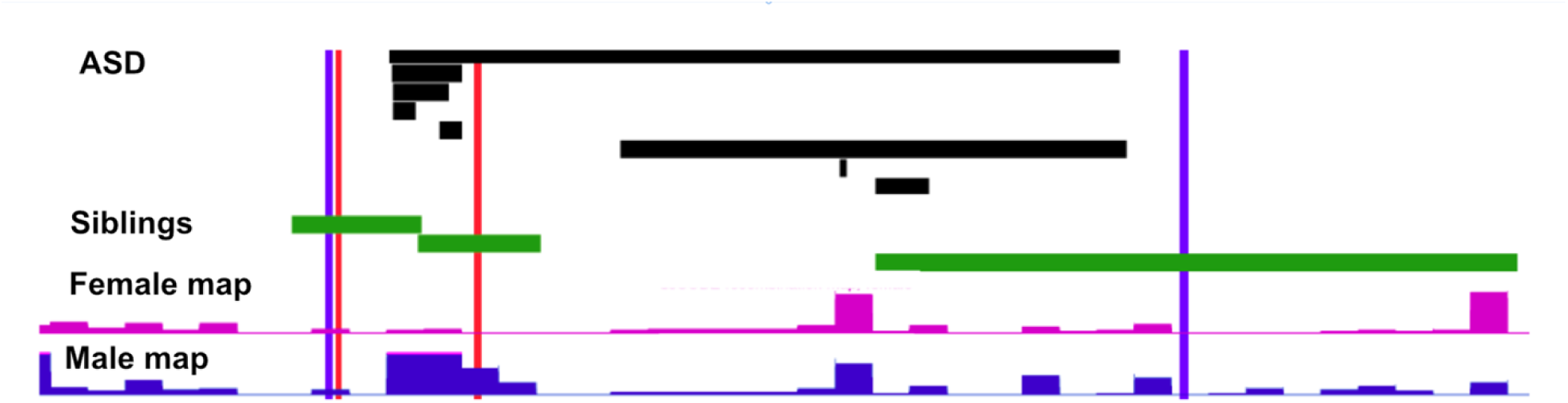
Comparison of recombination bin detection methods. An example of tracks of recombination intervals seen in ASD individuals (black) and their non-ASD siblings (green) for a chromosomal region. Decode recombination frequency map tracks are shown in pink for male and blue for female maps. The red vertical bars indicate the bounds of the bin defined by the minimum depth criterion, where there is a fully overlapping region for at least 4 ASD intervals, and a fifth is then included within a ± 15kb buffer. Blue vertical bars indicate the bounds defined by the cluster binning method, with 8 ASD intervals within 15kb of one another.

In evaluating the relative power of the methods, power improved in both methods when including a buffer region. The evaluation of the power of each method is described in detail in S1 Method. Based on the power analysis, a minimum recombination ‘depth’ of four recombination events was required to be considered a bin.

Two strategies for counting the number of intervals in each candidate region were also trialled. Both methods contrasted case and control recombination counts in the bin versus their counts in the rest of the same chromosome (27,28). In the “simple count Fisher’s exact test”, the number of recombinations that occurred in the candidate bin region was a simple count of any control recombination interval that overlapped the bin for any distance. In the “width weighted Fisher’s exact test” a sampling procedure accounted for the degree of certainty in the recombination location provided by the interval lengths (S2 Method). Briefly, for each recombination interval that overlaps the candidate bin, the algorithm selects a single point from a uniform distribution along the length of the interval. If the point is contained in the candidate bin, the interval is added to the appropriate case count or control count for that trial, and Fisher’s exact p-value is calculated using the sampled counts. The sampling process is repeated many times to obtain a vector **P** of Fisher’s exact test p-values. The final test statistic is the sample mean of the vector **P**. The sample mean was selected as the appropriate summary statistic due to the values of **P** being approximately normally distributed. The method is essentially a Monte Carlo estimate of the control recombinations in the candidate bin. The simple count method of all overlapping control intervals can be considered to be an upper bound of the width weighted Fisher’s exact test, in which all sampled points are contained in the candidate region.

### Permutation testing of fully informative families

In each of the proposed methods, the p-values generated by the test for enrichment are not immediately interpretable due to a highly non-standard underlying distribution, and thus can only be used as a means of ranking potential enriched regions (28). A permutation procedure was designed to control for structure in the family units, the distribution of recombination on individual chromosomes, and the underlying nonstandard distribution of the bin sizes (29–31). To account for the family structure of one affected child and at least two unaffected children, permutation involved in each replicate randomly selecting one child from each family to label as the affected child, and label all others as unaffected. Each permuted dataset was then analysed following the bin formation procedure of the main analysis to define maternal, paternal, and combined data. The proportion of permutations that had an enrichment at least as significant as that seen for the real data points were taken as the p-value. The permutation algorithm is described in detail in Supplementary Method S3.

### Evaluating recombination excess in partially informative pedigrees

Families having one affected child, one unaffected child and two parents genotyped are much more numerous in the dataset than those with at least two unaffected children. This structure permits the localisation of recombination events, but cannot determine which child carries the recombinant chromosome, thus these families are partially informative for recombination distribution. For these families, the recombination excess of certain genomic regions is not contrasted between case and controls from the same pedigree, rather by comparing the excess of recombination events across both proband and unaffected with the expectation from an external recombination map (32).

Two candidate methods were evaluated for power: “narrow bins” encompassing the region of maximum recombination interval coverage at a single point, with a 15kb buffer on each side; and “wide bins” were defined as the maximum extent of any recombination intervals that overlap the point of maximal recombination interval coverage. Intervals of longer than 200kb were filtered in quality control, so that the total width of a wide bin will not significantly exceed 400kb. For both investigations, the width weighted Fisher’s exact test p-value was estimated, as it was determined as optimal in the three child families. A contingency table illustrating the components of the test is available in Supplementary Method S4. A resampling-based power calculation was necessary to determine the more powerful approach, as the effect of the expected number of recombinations derived from the recombination rate maps on the variance of the test for enrichment was uncertain. Furthermore, the absence of case/control labels given the use of the map as a reference required the use of a bootstrap-based method rather than the permutation method that was possible in the fully informative pedigrees. Descriptions of the bootstrap method and corresponding power calculation is available in S5 and S6 Methods. The narrow binning method was slightly more powerful than the wide binning method, with 10% power to detect enrichment at an effect size defined by a nominal p-value *ϕ* = 1 × 10^−11^ and 100% at *ϕ* = 1 × 10^−15^. In contrast, the wide binning method had 10% power at *ϕ* = 1 × 10^−14^, 70% power at *ϕ* = 1 × 10^−15^ and 100% at *ϕ* = 1 × 10^−16^. While the difference in power is slight and the effect sizes required for non-zero power to detect a genome-wide significant locus are extremely high, narrow binning is the more powerful method and is therefore used as the method to detect enrichment.

The minimum criterion to define a bin was 30 recombinations, since this bin size had reasonable power to detect a recombination excess over a range of effect sizes (S1 Fig).

## RESULTS

### Fully informative pedigree recombination distributions are similar in autistic children and non-autistic siblings

The gross pattern of recombination appeared similar among autistic children and their non-autistic siblings in the fully informative (3+ child) families. There was no excess in total recombination among autistic children versus non-autistic siblings for any chromosome or for the genome as a whole (whole genome p = 0.24) (S1 Table) (33). There was no excess recombination in autistic children in the centromeric region for any chromosome or the whole genome (whole genome p = 0.94) (S2 Table) (27,28). There was no excess recombination in autistic children in coldspots, defined as a region of 50kb or longer with a cM/Mb ratio of 0.5 or less (whole genome p = 0.20) (S3 Table) (14). Runs of homozygosity are associated with autism in both multiplex and simplex cases (34,35). However, Kolmogorov-Smirnov tests found that there was no systematic difference in recombination event interval lengths (in base pairs) between cases and controls (combined p = 0.32; maternal p = 0.82; paternal p = 0.29). Among maternal recombinations, 9.1% (980/10,607) were *>*200kb in autistic children compared to 9.2% (2,057/22,630) in the non-autistic children (p = 0.68). Among the paternal recombinations, 9.0% (621/6,909) were *>*200kb in autistic children compared to 8.1% (1,198/14,839) in non-autistic children (p = 0.04; combined p = 0.11, S2 Fig).

### Regions enriched for *de novo* recombination in fully informative pedigrees

The minimum depth recombination region detection algorithm identified 117 maternal, 99 paternal, and 472 combined recombination bins of 4 or more case recombination events with an odds ratio of case to control recombinations of greater than one. Of those, 31 combined loci, 13 maternal loci, and seven paternal loci had a nominal width-weighted Fisher’s exact p-value that was less than 0.05 (For comparison, there were 25 combined, two maternal, and seven paternal regions that met the minimum case-count threshold but whose odds ratio of case to control recombinations was less than one).

After applying the permutation procedure, none of the regions remained genome-wide significant (Table 1). However, there were six loci that had especially low p-values and merited further examination (Fig 3, Table 1), as well as another thirteen loci identified in the maternal and paternal recombination sub-analyses (Fig 3, S5 Table). A comprehensive analysis of copy number variation in Simons Simplex Collection did not report any copy number variation, either duplication or deletion, in any of the recombinant probands in any of the nineteen recombination regions identified in this study (36).

**Fig 3.**
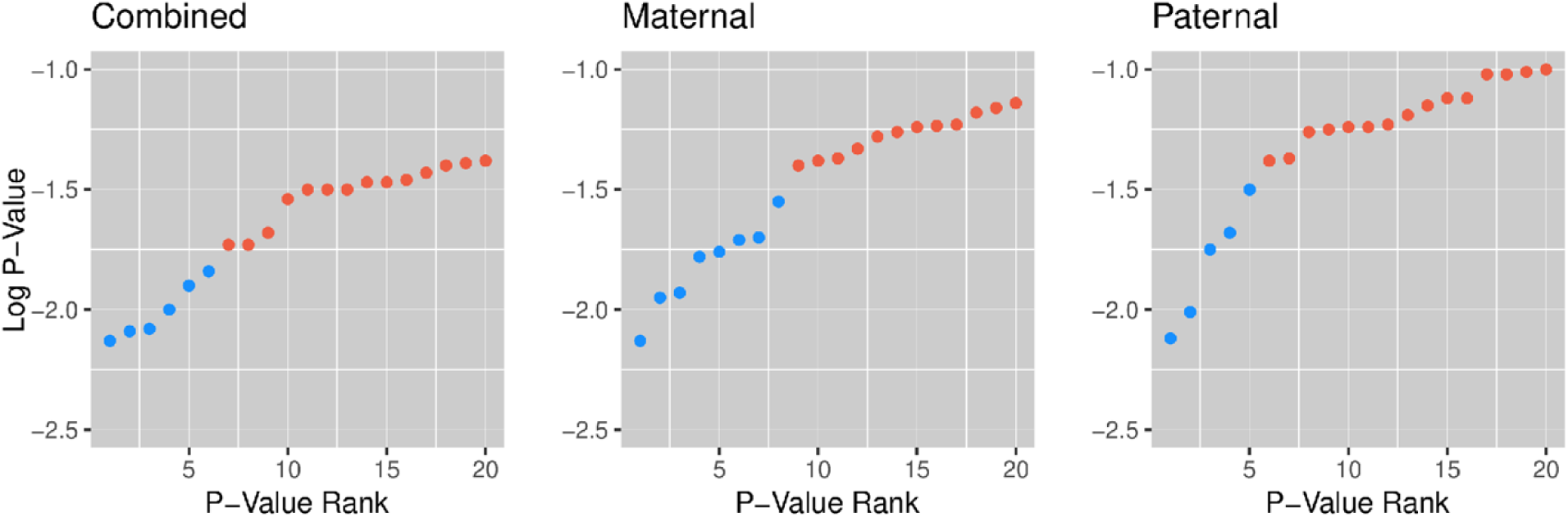
Distribution of low p-value regions in each parental grouping. The ‘elbow’ is a visual heuristic to identify regions for further investigation rather than a rigorous test of significance. Six combined, eight maternal, and five paternal regions fell below the elbow and were investigated further for gene content and potential mechanisms of contribution to autism.

**Table 1.** Loci with suggestive levels of enrichment for combined maternal and paternal recombination.

| **Fully Informative Pedigrees (three-child families)** |  |  |  |  |  |  |
| --- | --- | --- | --- | --- | --- | --- |
| Chromosome Band | Start (MB) | End (MB) | Case Count | Control Count | Width-Weighted P-Value | Permuted P-Value |
| 20q13.33 | 59.537 | 59.575 | 9 | 7 | 0.0072 | 0.2570 |
| 15q11* | 20.066 | 20.175 | 7 | 2 | 0.0082 | 0.2863 |
| 16q23.1 | 79.116 | 79.151 | 8 | 2 | 0.0086 | 0.2966 |
| 6p21.2 | 37.557 | 37.590 | 5 | 1 | 0.0098 | 0.3260 |
| 5p15.32* | 5.193 | 5.225 | 5 | 2 | 0.0097 | 0.3879 |
| 14q22.3 | 56.365 | 56.397 | 4 | 1 | 0.0138 | 0.4133 |
| **Partially Informative Pedigrees (two-child families)** |  |  |  |  |  |  |
| Chromosome Band | Start (MB) | End (MB) | Observed Count | Expected Count | Width-Weighted P-Value | Bootstrap P-Value |
| 19p13.2* | 7.228 | 7.325 | 63 | 1 | 6.98E-09 | 0.9810 |
| 11q13.3 | 69.698 | 69.807 | 32 | 1 | 1.55E-07 | 1.0000 |
| 10q26.13* | 125.853 | 125.935 | 44 | 3 | 1.66E-07 | 1.0000 |
| 10p15.3* | 1.502 | 1.534 | 87 | 24 | 1.82E-07 | 1.0000 |
| 2q14.3 | 129.010 | 129.042 | 52 | 10 | 7.92E-06 | 1.0000 |
| 10q11.23 | 51.579 | 51.800 | 33 | 7 | 2.21E-05 | 1.0000 |
| 9q34.3 | 139.151 | 139.233 | 42 | 6 | 2.90E-05 | 1.0000 |

### Regions enriched for *de novo* recombination in partially informative pedigrees

Among the two-child families, there were 566 combined loci (overlapping in large part with 103 maternal-recombination only loci and 107 paternal loci) that met the minimum depth requirement of 30 recombinations at a single point. 34 combined loci had a nominal width-weighted Fisher’s exact p-value of less than .05. After correction for multiple testing, none of the regions remained genome-wide significant given the threshold for genome-wide significance derived from the bootstrapping procedure at 3.9×10^−15^ for combined data (1.8×10^−8^ for maternal and 3.0×10^−14^ for paternal). The very tiny thresholds defined from the bootstrap could reflect a small set of regions with consistent deviations between the SSC cohort and the deCODE reference genetic maps, which have slightly different ranges, owing to the expansion over time of genome assembly GRCh37 to which both correspond. However, seven combined loci had especially low p-values warranting further investigation (Fig 4), with none of these located in centromeric or telomeric regions most prone to differences among genetic maps (Table 1, S4 Table, S5 Table). As in the case of the three-child families, no copy number variation was detected in either child in the families with recombinations identified in these regions (36).

**Fig 4.**
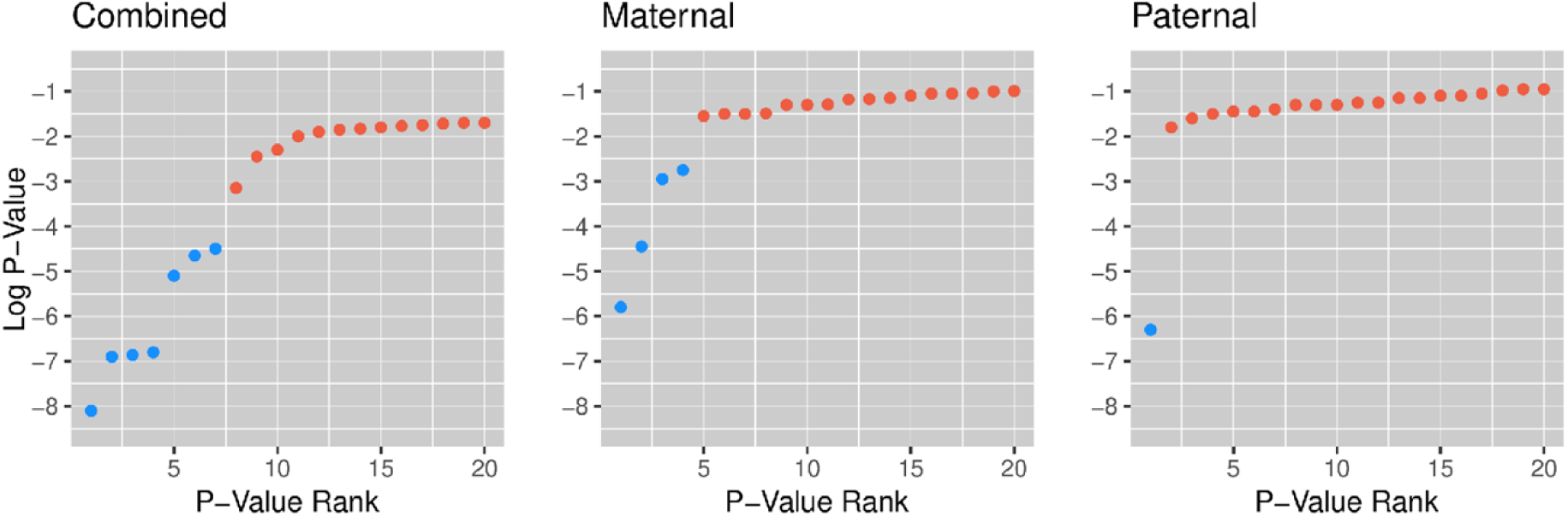
Width-weighted Fisher’s exact p-values for recombination hotspots detected in partially informative families. Seven combined loci had especially low nominal p-values and merited further investigation. Blue points were investigated further, while red points were not.

### Enrichment testing of loci identified in fully informative families within the partially informative families

.A modified genetic map-based enrichment test was used to examine whether any of the intervals of interest from fully informative families (Table 3) was enriched for recombination in the families with two children. For each locus, the expected number of recombinations in the hotspot was computed using the genetic map recombination rate. All recombination intervals that overlapped the bounds of the recombination hotspot were identified and used to compute the width-weighted Fisher’s exact test for the region in the two-child families (S6 Table). All but one of the seventeen loci that were within the map ranges had an excess of recombination as compared to the genetic map, as indicated by odds ratios of greater than one. The lowest nominal p-value (nominal width-weighted Fisher’s exact p-value = 0.009) occurred in the combined 20q13.33 locus. This interval also showed the most significant p-value in the combined data in the families with three or more children (Table 3.6), suggesting that it may indeed be a genuine association. None of the other loci had a nominal width-weighted p-value that was lower than 0.05.

### Location of recombination hotspots with respect to nearby protein-coding genes

The thirteen regions of notable recombination excess identified during the parental combined analysis of both two-child and three-child cohorts included five regions where the interval of recombination only overlapped a single protein-coding gene, and seven where the interval lay in between two protein-coding genes (S4 Table). One interval (at 15q11) occurred next to the centromere, a region where protein coding genes are sparse, and where the nearest downstream putative protein coding gene was *GOLGA6*, a member of a gene family implicated in genetic rearrangement on chromosome 15 (37–42); another interval (10q11.23) spanned seven genes, making it hard to pinpoint which genes might be implicated in ASD. Leaving out these two regions, we investigated whether these genes were included in the 7,501 high confidence and candidate SFARI gene database genes listed on the Genetrek resource (43). The five overlapped protein-coding genes (*WWOX, ADAMTS16, INSR, ADARB2, and HS6ST1)* were identified as high confidence or candidate ASD genes (p=0.02, Fisher’s exact test).

Among the twelve protein-coding genes that flanked intervals that did not directly overlap protein coding genes (*ARHGEF18, ZNF557, WDR37, PFKP, UGGT1, RAB6C, CDH26, CDH4, CMTR1, MDGA1, KTN1, PELI2)* there was no particular enrichment for candidate and high confidence ASD genes (OR=1.13, p=1). Taking the five loci that overlap a single protein coding gene, together with the cadherin locus confirmed in the two-child resource, these six candidate loci represent useful candidate genes to consider further.

### Potential roles of the six most implicated genes

S3 Fig illustrates the recombination distributions observed at the 20q13.3 region, with enrichment between two genes that encode two cell adhesion cadherin proteins (CDH4 and CDH26). Some identical interval termini are seen in different recombination events, which is likely to arise from the haplotype structure yielding marked boundaries between more informative and less informative regions. Seven other cadherin proteins and 20 protocadherins have been associated with ASD (44–46). *CDH26* showed a weak association with Tourette Syndrome (47). *A* child with Landau-Kleffner Syndrome, a condition which causes seizures and speech difficulties, who inherited an intronic deletion in *CDH4* from their father who had experienced a transient regression in speech during childhood (48). *CDH4* is listed in the SysNDD curated database of candidate neurodevelopmental disorder (NDD) genes, with two rare coding mutations identified in the gene in children with NDDs (49). Cadherins appear to affect both the development of a range of brain tissues and the mature function of synapses. S4 Fig illustrates that there is little typical population linkage disequilibrium in this region. The combination of variants showing the greatest disequilibrium on opposite sides of the hotspot (and therefore capable of disruption by recombination) showed no significant association with ASD in the cohort (S7 Table). Thus, any effect is unlikely to be mediated by recombination among common variants, and may reflect involvement of rare variants or rearrangements that do not result in changes in copy number.

*WWOX* (Fig 5) is a very large gene (1.11Mb) at the 16q23.1 locus that can act as a tumour-suppressor gene (50). *WWOX* has been linked with rare variants and copy number variation in multiple autism probands in the Simons Simplex Collection (43,51). *WWOX* contains the genomic instability hotspot *FRA16D*, the second most common chromosomal fragile site. *FRA16D* mediates copy number variation and other rearrangements (50). Loss of function mutations in *WWOX* are associated with abnormalities in DNA damage repair. While no marked excess of copy number variation was seen at this locus in the Simons Simplex Collection cohort, loss of function point mutations and copy number variation have been associated with autism spectrum disorder, in both intragenic variants and larger rearrangements in the region (36,52). While recombination may be implicated in de novo rearrangements, it could also generate novel haplotypes within *WWOX* itself by combining genetic variants in the last protein coding exon or 3’UTR with other variants upstream. The recombination prone region of slow DNA replication in *WWOX* peaks at intron 5, *WWOX* is also known to form translocations with chromosome 14q32 immunoglobin heavy chain (*IGH*) in multiple myeloma (50). Nearby genes upstream *CLEC3A* (53,54) and downstream *MAF* are also implicated in neurodevelopmental disorders (43,55). It is possible that complex recombinations involving *WWOX* and *MAF* could generate risk haplotypes: since these two genes appear to act together in cancer, they may also act together in ASD (50). Mouse *WWOX* has been identified as one of a very small number of genes undergoing double strand breaks and genomic rearrangements in somatic tissues (56,57), suggesting the hypothesis that somatic recombination may even contribute to brain cell diversity (57).

**Fig 5.**
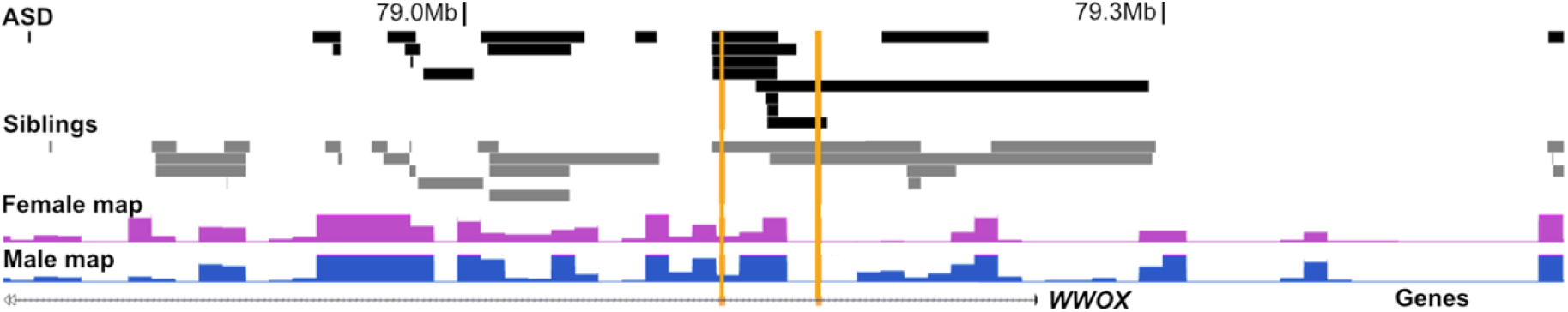
Recombination events in *WWOX*. Map of individual recombination event intervals in the 16q23.1 region of excess ASD recombination (bounded by vertical orange lines) seen in three child families within the last intron of the *WWOX* gene (black: ASD; grey: non-ASD siblings). Note: controls have greater than double the sample size of cases.

The 5p15.32 recombination locus overlaps *ADAMTS16* (Fig 6), a disintegrin-like and metalloproteinase with thrombospondin type 1 motif 16, associated with Cri-du-Chat Syndrome, a rare neurodevelopmental disorder resulting from a deletion of variable size in 5p (58). Cri-du-chat, often caused by the de novo deletion of the 10-20Mb region at the distal end of the 5p arm, involves feeding difficulties, hypotonia, characteristic facial features, behavioural problems including hyperactivity and repetitive movements, and difficulties with speech (59). The recombination interval spans three internal exons, so novel haplotypes may disrupt its function. Depletion of the nearby gene *ICE1* leads to increased abundance of nonsense-mediated decay targets in other autism genes (60,61).

**Fig 6.**
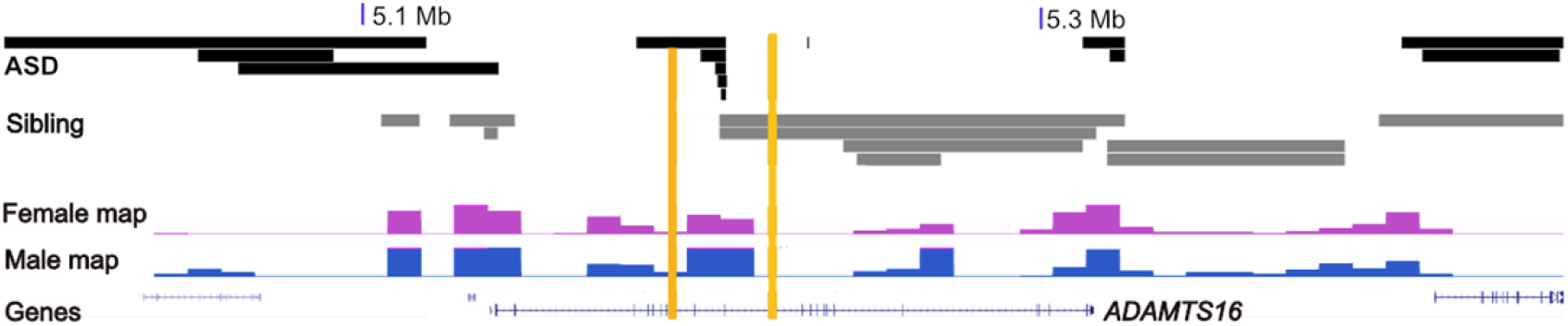
Recombination events in *ADAMTS16*. Recombination events in a region of excess ASD recombination (bounded by vertical orange lines) in three-child families in 5p15.32.

Recombination excess in the 19p13.2 band overlapping *INSR* was enriched in the partially informative (2-child) families (Fig. 7). *INSR* encodes Insulin receptor, a member of the receptor tyrosine kinase family of proteins (62). Mutations in this gene are associated with severe insulin resistance (63). There is no evidence in the literature of an association between *INSR* and autism spectrum disorder.

**Fig 7.**
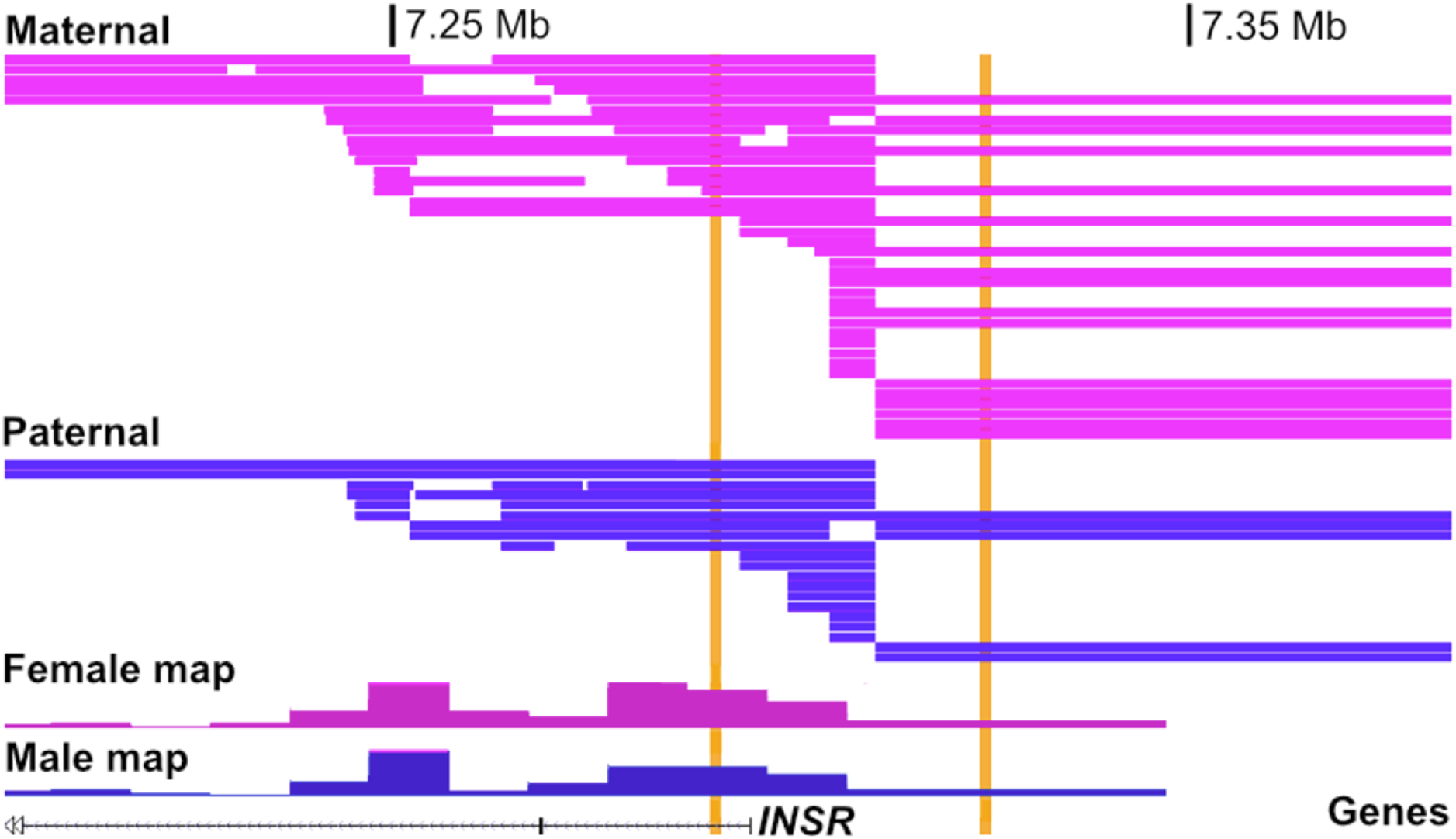
Recombination events overlapping *INSR*. A region of recombination excess (bounded by vertical orange lines) relative to the reference recombination map, seen in partially informative two-child families in the 19p13.2 region. Maternal recombinations are shown in pink and paternal recombinations are shown in blue.

The enriched interval in the10p15.3 band overlapped *ADARB2* (Fig. 8). *ADARB2* encodes Adenosine deaminase RNA specific B2. At least 21 cases of 10p15.3 microdeletion have been reported, with patients showing varying degrees of developmental disability and speech delay (64). A further study identified monoallelic expression in *ADARB2* in the first intron in individuals with ASD (65). The nearest proximal gene to the recombination bin is *WDR37*, which encodes a member of the WD repeat protein family, a large family present in all eukaryotes that functions in a wide variety of cellular processes. *WDR37* is the only gene identified in the recombination hotspots that is included in the GeneTrek high confidence NDD candidate gene list. There are numerous reports of *WDR37 de novo* missense mutations resulting in developmental delay, intellectual disability, and epilepsy (66).

**Fig 8.**
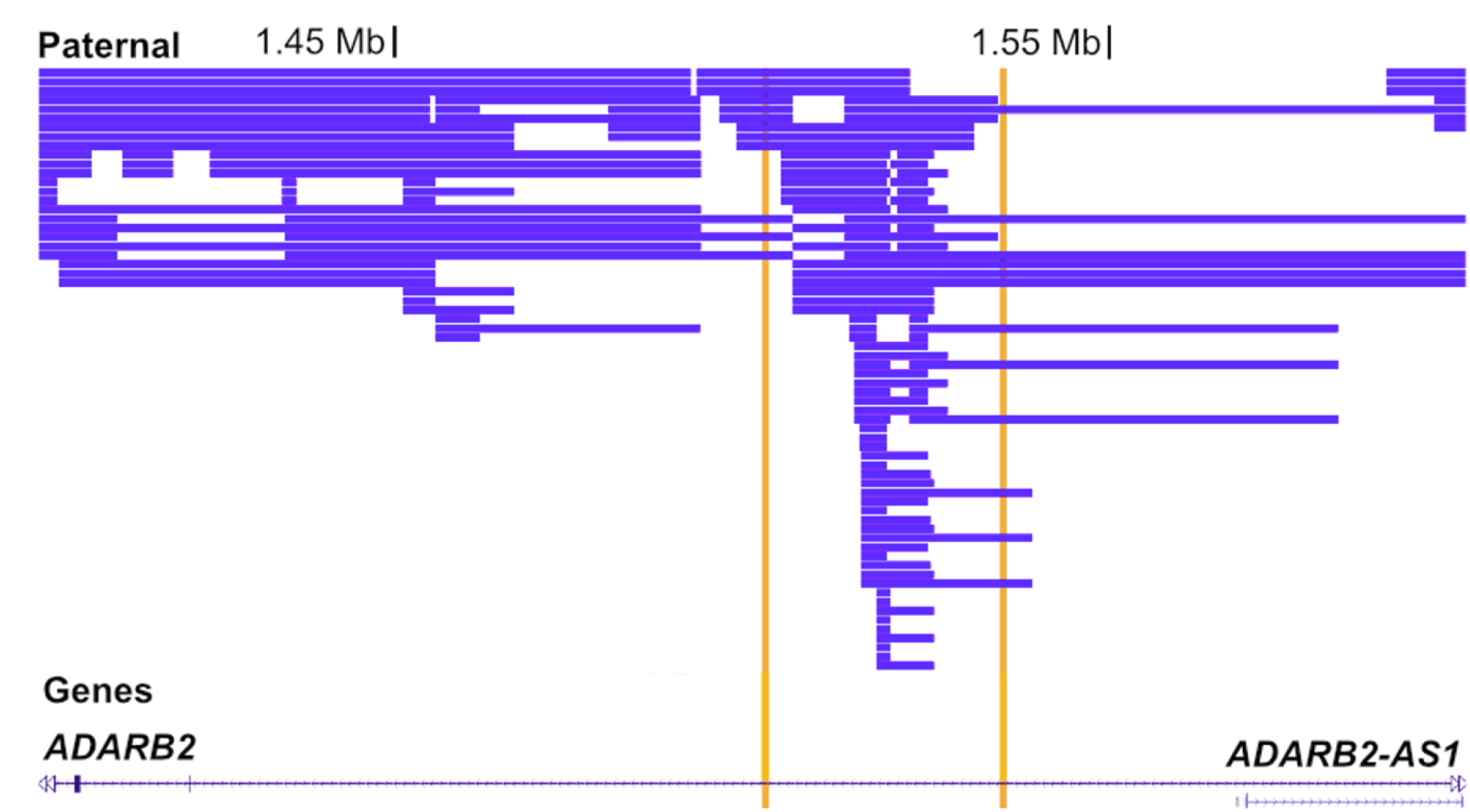
Recombination events in *ADARB2*. Recombinations in a region of recombination excess (bounded by vertical orange lines), all paternal, in partially informative two-child families in 10p15.3. The sub-telomeric region is outside the range of deCODE genetic maps, but is within the range of the maps published by Bhérer, *et al* (28). No maternal recombination events were seen in this region.

The recombination-enriched interval at 2q14.3 overlapped *HS6ST1* (Fig. 9). *HS6ST1* encodes Heparan sulfate 6-O-sulfotransferase 1, which is involved in extracellular sugar modifications (67). *HS6ST1* is included in the GeneTrek candidate NDD gene list on the basis of a high probability of being loss-of-function intolerant (pLI *>*0.9) and being associated with epilepsy in one or more of the GeneTrek constituent epilepsy gene databases (43,68).

**Fig 9.**
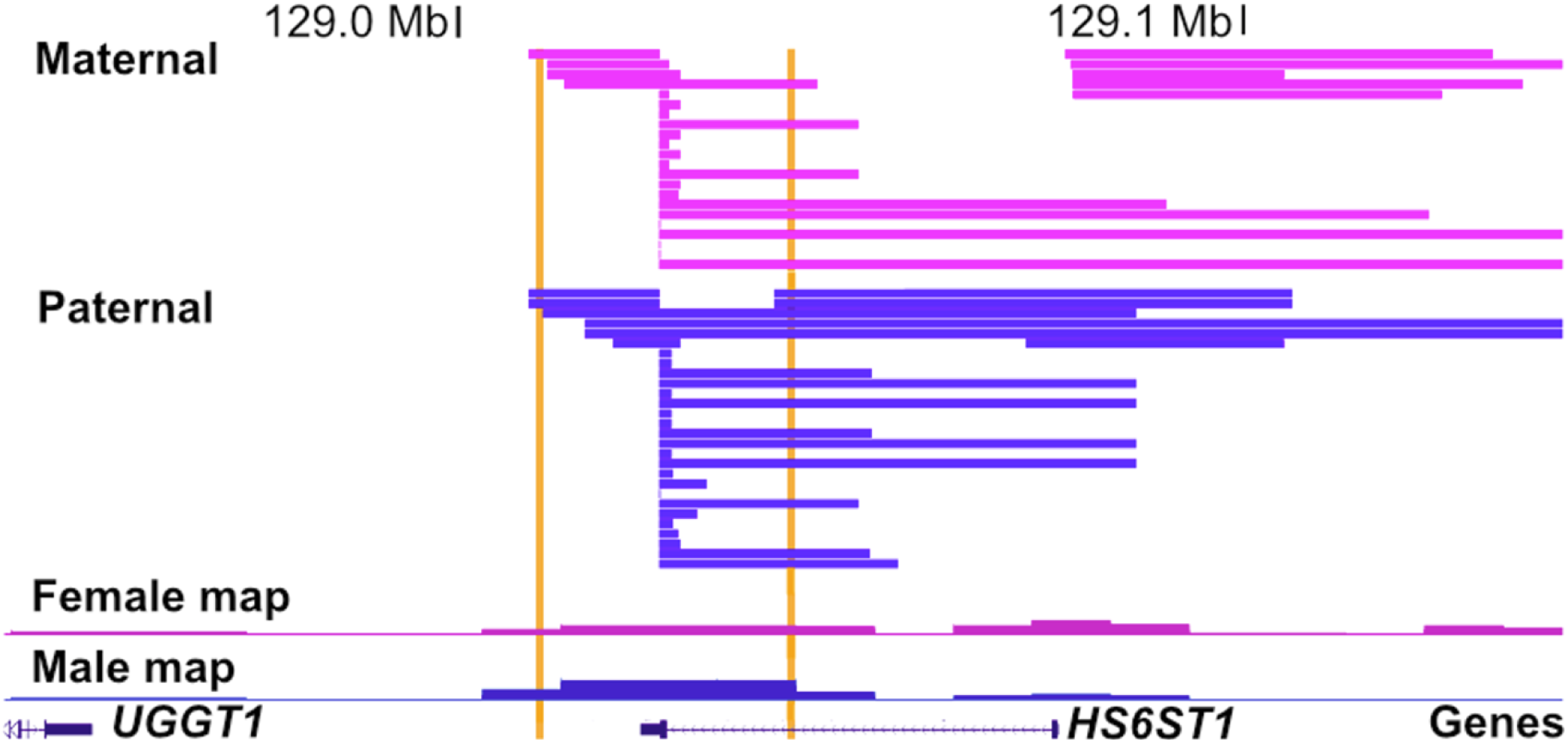
Recombination events overlapping *HS6ST1*. Recombinations in a 2q14.3 region of recombination excess (bounded by horizontal orange lines) in partially informative two-child families (pink: maternal; blue: paternal).

While the association of *HS6ST1* with ASD has not been explored in depth, heparan sulfate is ubiquitous in the regulation of cellular processes during embryonic and neonatal brain development, and many regulatory disruptions that result in a deficit of heparan sulfate are associated with severe neurodevelopmental disability, making *HS6ST1* a plausible novel autism candidate gene (69).

For the three loci from the fully informative families (*WWOX, ADAMTS16 and CDH26-CDH4* regions), we investigated whether there was any association of ASD behavioural phenotypes with recombination events in that region. Behavioural data for the Simons Simplex Collection were recorded using the Aberrant Behavior Checklist (ABC), a 58-question checklist of non-adaptive behaviours related to autism.

Responses are typically scored by a parent and occasionally by a professional on a scale from 1-4, with a higher score indicating a greater severity of the symptom. The checklist is divided into five subscales, titled (1) Irritability, Agitation, Crying (15 items); (2) Lethargy/Social Withdrawal (16 items); (3) Stereotypic Behavior (7 items); (4) Hyperactivity/Noncompliance (16 items); and (5) Inappropriate Speech (4 items). Evaluation by totaling the score across the five subscales is explicitly discouraged in the manual (70,71). While there was a weak suggestion that recombination in *WWOX* was associated with lower hyperactivity, the effect is not significant after Bonferroni correction for multiple testing (Fig. 10, Table 2).

**Fig 10.**
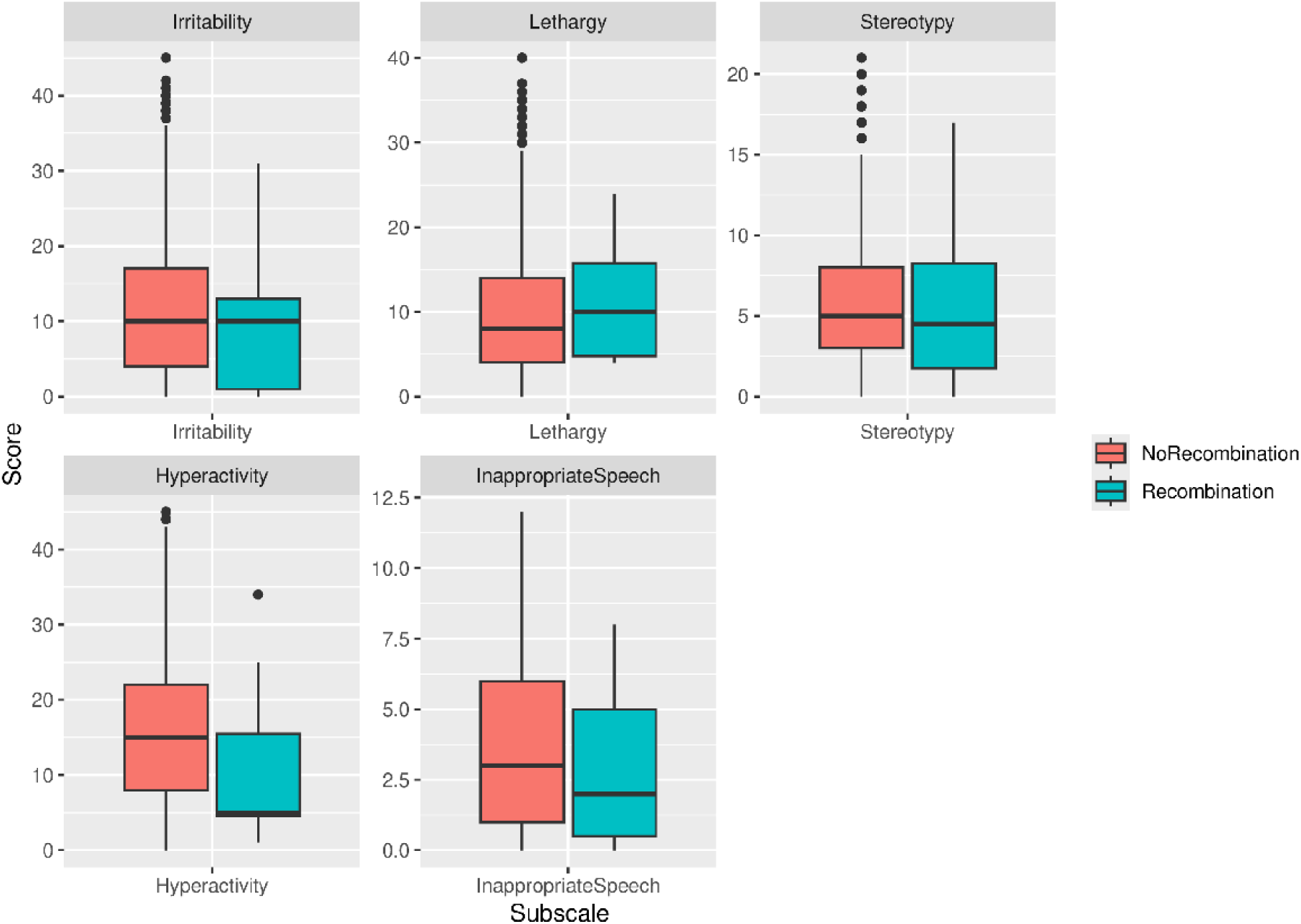
Phenotype comparison of Aberrant Behavior Checklist phenotype subscales. ABC phenotype score comparisons at the 16q23.1, where there is a weak suggestion of higher hyperactivity scores among probands with a recombination (red) at the locus versus those without (blue).

**Table 2.** Kolmogorov-Smirnov p-values for recombinant vs non-recombinant proband behavioural phenotypes for three-child families. After Bonferroni correction, there is no significant difference between the phenotype distributions in the recombinant and nonrecombinant probands at any particular locus.

| Band | Gene | Irritability | Lethargy | Stereotypy | Hyperactivity | Inappropriate Speech |
| --- | --- | --- | --- | --- | --- | --- |
| 20q13.33 | CDH26-CDH4 | 0.48 | 0.47 | 0.42 | 0.90 | 0.08 |
| 16q23.1 | WWOX | 0.35 | 0.78 | 0.87 | 0.04 | 0.91 |
| 6p21.2 | ADAMTS16 | 0.89 | 0.72 | 0.16 | 0.29 | 0.12 |

## DISCUSSION

This study defines a set of methods suitable for establishing regions of excess recombination among a cohort of patients, and applies it to the Simons Simplex Collection (SSC) of families with one affected child with ASD and one or more siblings additionally genotyped along with both parents. The enrichment of neurodevelopmentally associated genes in the five implicated loci where the recombination interval only overlapped a single protein coding gene is suggestive that there are indeed possible excesses of recombination leading to increased risk of ASD.

Such an excess of recombination may be mediated by association with damaging non-reciprocal rearrangements, such as copy number changes, with inversions or translocations that alter the regulatory environment of genes. These may involve homologous recombination of highly similar sequences of at least 300 bp, and in the case of tandem repeats, typically >1kb in length with >95% sequence identity or non-homologous recombination involving only small regions of sequence similarity and often much more complex rearrangements that likely reflects an attempt to repair DNA damage (72). The common copy number variants (CNVs) typically arise through homologous recombination, and the loci we identified here seem unlikely to reflect enrichment for such *de novo* events, since none of them were identified as CNV enriched in a survey of this cohort of families, and no CNVs were identified in the recombinant individuals. In the case of *WWOX*, this locus has exceptionally large introns, and is implicated in recombinational repair. For the other loci, there may well be a role of recombination that is not only strictly homologous, but entirely reciprocal, yielding unfavourable haplotypic combinations that have an effect on neurodevelopment. Such recombinations could arise between adjacent genes of the same family, who may have maintained common mechanisms of expression control via long-range enhancers, such as the CDH4-CDH26 region highlighted here, although our haplotype analysis at this locus suggests that such combinations do not appear to be generated from recombination only between common variants, and are therefore more likely to involve some rare variants. Recombinations among protein coding or gene controlling variants within the gene itself may therefore play a role. Extensive sequencing of candidate loci is required to provide further evidence, ideally in a separate cohort to this discovery cohort.

The study here identifies a potential role for recombination in neurodevelopmental disorders. However, the low level of replication between the two child and the three child cohorts indicates that if there is a signal, there is low power to detect it. Thus, very large sample sizes are likely to be needed. It may well be that long range sequencing among larger cohorts will identify such rare recombination generated variants. In such a study design, a single proband and two parents are sufficient to provide information, and so the available cohort sizes may be correspondingly larger. An integration of the two-child and three-child recombination mapping study designs used here, alongside sequencing information, is the likely best approach for defining recombination as a cause of de novo ASD. With such approaches in place, it may even be feasible at some point in the future to incorporate some of this information into genetic counselling for parents of a child, since the identification of a recombination event in an implicated locus could feasibly provide some evidence of a potential *de novo* causation that might influence their family planning considerations, in the absence of any other information regarding whether the child’s ASD is likely arising from inherited or from *de novo* elements.

## Data Availability

The genetic data used in this study were obtained from the Simons Simplex Collection (SSC) via SFARI Base (https://www.sfari.org/resource/sfari-base/). These data contain personally identifiable information from human research participants and cannot be made publicly available by the authors under the terms of the data access agreement and participant consent. Approved researchers can obtain the SSC population dataset described in this study by applying at https://base.sfari.org.

The inferRecom software is available at https://github.com/catherinefayemahoney/inferRecom.

## Acknowledgements

Data for this study were provided by the Simons Foundation Autism Research Initiative. We are grateful to all of the families at the participating Simons Simplex Collection (SSC) sites, as well as the principal investigators (A. Beaudet, R. Bernier, J. Constantino, E. Cook, E. Fombonne, D. Geschwind, R. Goin-Kochel, E. Hanson, D. Grice, A. Klin, D. Ledbetter, C. Lord, C. Martin, D. Martin, R. Maxim, J. Miles, O. Ousley, K. Pelphrey, B. Peterson, J. Piggot, C. Saulnier, M. State, W. Stone, J. Sutcliffe, C. Walsh, Z. Warren, E. Wijsman). We appreciate obtaining access to phenotypic and genetic data on SFARI Base.

## SUPPORTING INFORMATION CAPTIONS

**S1 Method. Power calculation for candidate enrichment detection methods in fully informative pedigrees.**

**S2 Method. Width-weighted Fisher’s exact test.**

**S3 Method. Permutation procedure for fully informative pedigrees**

**S4 Method. Contingency table for width-weighted Fisher’s exact test in fully informative pedigrees.**

**S5 Method. Single-step p-value adjustment for partially informative pedigrees.**

**S6 Method. Power calculation for narrow and wide binning methods.**

**S1 Fig. Power to detect enrichment by minimum bin depth.** Power to detect enrichment in a single bin. Each curve represents a bin of the given size. The improvement from ten to twenty and subsequently twenty to thirty is very large, while after thirty the improvement in each bin depth decreases. On this basis, a minimum criterion of 30 recombinations was selected as the criterion for a putatively enriched region in partially informative pedigree data.

**S2 Fig. Distribution of interval widths for maternal, paternal, and combined intervals.** In each case, the distribution of lengths is highly similar between autistic and non-autistic children (Kolmogorov-Smirnov p-values: combined p = 0.3211; maternal p = 0.8249; paternal p = 0.2945). There is a nominally significant (p = 0.0387) excess of long (*>*200kb) paternal intervals in autistic children, which did not remain significant after Bonferroni correction.

**S3 Fig. Combined 20q13.33 Locus.** The locus lies between two cadherins, CDH4 and CDH26. While it is out of the range of the deCODE recombination rate maps in the browser, it is within the range of the more recent recombination rate maps produced by Bhérer, et al.

**S4 Fig. Linkage disequilibrium in the 100kb region centred on the 20q13.33 recombination hotspot.** The pictured region lies 915kb distal to CDH26 and 222kb proximal to CDH4. The lower triangle shows linkage disequilibrium between SNPs computed using r^2^, with higher r^2^ values pictured in dark red. The upper stacked bar chart shows the number of recombination intervals beginning (blue) and ending (red) at that SNP. The endpoints of the recombination hotspot are indicated by black arrows. There does not appear to be strong linkage disequilibrium between any SNPs on opposite sides of the hotspot.

**S1 Table. Recombination rate Mann-Whitney U-test p-values.** Mann-Whitney U-Test pvalues for stochastic difference in recombination rate between autistic and non-autistic children. After Bonferroni correction, none of the maternal, paternal, or combined chromosomes had a significant difference in recombination rate between autistic and non-autistic children (combined whole genome p = 0.244).

**S2 Table. Fisher’s exact test p-values for recombination rate near the centromere.** The centromeric region was defined as a 3Mb region about the centromere, with coordinates obtained from genome assembly GRCh37 (hg19). After Bonferroni correction, no maternal, paternal, or combined chromosome or genome had a significant difference in total recombination rate near the centromere in autistic children versus unaffected siblings.

**S3 Table. Fisher’s exact test p-values for recombination rate in coldspots.** The cold spots were defined as a region of at least 50kb with a recombination rate of less than 0.5cM/Mb. After Bonferroni correction, no maternal, paternal, or combined chromosome or genome had a significant difference in total recombination rate in cold spots in autistic children versus unaffected siblings.

**S4 Table. Suggestively enriched loci in combined data with nearby genes.**

**S5 Table. Suggestively enriched loci in parental subgroups.**

**S6 Table. Test for enrichment versus genetic map in loci identified in fully informative pedigrees.** One locus in the chromosome 15 centromere region could not be tested as it was out of the range of the genetic maps. The lowest width-weighted Fisher’s exact p-value occurred in the combined 20q13.33 region.

**S7 Table. Genotype combination frequencies for autistic and non-autistic children at rs6128970 and rs77396656.** There is no significant difference in the distribution of genotype combinations between the two groups (Chi-Sq p = 0.9565)

